# A starting guide on multi-omic single-cell data joint analysis: basic practices and results

**DOI:** 10.1101/2024.03.30.587427

**Authors:** Lorenzo Martini, Roberta Bardini, Stefano Di Carlo

## Abstract

Multi-omics single-cell data represent an excellent opportunity to investigate biological complexity in general and generate new insights into the biological complexity of heterogeneous multicellular populations. Considering one omics pool at a time captures partial cellular states, while combining data from different omics collections allows for a better reconstruction of the intricacies of cell regulations at a particular time. However, multi-omics data provide only an opportunity. Computational approaches can leverage such opportunities, given that they raise the challenge of consistent data integration and multi-omics analysis. This work showcases a bioinformatic workflow combining existing methods and packages to analyze transcriptomic and epigenomic single-cell data separately and jointly, generating a new, more complete understanding of cellular heterogeneity.

## 1 Introduction

Cellular heterogeneity is a fundamental characteristic of multicellular organisms that arises during their developmental processes. The existence of diverse cell populations is essential to maintain homeostasis and coordinate physiological activities within an organism [7, 18, 2, 24, 28]. Each distinct cellular phenotype possesses a unique set of functional and structural features. Investigation of cellular heterogeneity contributes to our understanding of the structural composition of multicellular organisms by uncovering subtypes of cells that share a general cell type but exhibit different functional characteristics [5]. By studying heterogeneity, we can gain insight into physiological and pathological conditions that affect individual cellular phenotypes and the overall cellular composition of affected tissues [4].

To study cellular heterogeneity, single-cell Next Generation Sequencing (NGS) technologies have been developed, enabling high-throughput analysis of various biological signals at the single-cell level. These technologies encompass multiple omics pools, including transcriptomic (the most prevalent), proteomic, and epigenomic states of cells [40, 17]. Leveraging these technologies allows for the characterization of cell states [39, 22] and provides a deeper understanding of cellular regulatory processes by examining correlations between different omics.

In recent years, the emergence of multi-omics single-cell technologies has revolutionized the field by enabling the simultaneous profiling of multiple omics on the same set of cells [8]. These advancements have provided unprecedented insights into cell states and are poised to propel further research in this area. Several combinations of multi-omics techniques have been developed, and a particularly promising approach for studying cellular heterogeneity involves integrating single-cell RNA sequencing (scRNA-seq), which captures transcriptional expression levels, with single-cell assays for transposase-accessible chromatin sequencing (scATAC-seq), which measures genome-wide chromatin accessibility.

This paper presents a comprehensive yet simpe bioinformatic procedure for integrating scRNA-seq and scATAC-seq datasets to study cellular heterogeneity. It specifically focuses on best practices, reviews available pipelines for analyzing transcriptomic and epigenomic data, and discusses tools for correlating and jointly analyzing these datasets. Finally, the paper provides a practical example of applying the proposed procedure to investigate cellular heterogeneity in a multi-omics dataset of human peripheral blood mononuclear cellss (PBMCs) [1].

## 2 MATERIALS AND METHODS

Single-cell technologies have matured and numerous processing pipelines have been developed [13]. Notably, the processing of scRNA-seq data has become well-established and widely adopted [19, 12]. On the contrary, the analysis of epigenomic data, particularly scATAC-seq data, requires further investigation [6].

A lack of dedicated pipelines or established procedures exists regarding multi-omics data analysis. Researchers often resort to pipelines designed for specific omics (such as scRNA-seq or scATAC-seq) without clear guidelines on how to combine the results into a comprehensive multi-omics analysis.

To address the need for a combined analysis of scRNA-seq and scATAC-seq data, it is beneficial to utilize pipelines that provide functionalities for both data types. Three well-known tools in the single-cell field are particularly suitable: Monocle3 [35], Seurat [32], and Scanpy [38]. Although originally developed for scRNA-seq analysis, these tools offer companion packages for analyzing scATAC-seq datasets, namely Cicero [27], Signac [33], and Episcanpy [10] (Figure 1 (B)). An advantage of using these coupled packages is their shared underlying algorithms, facilitating the integration of results and mitigating the introduction of artifacts resulting from incompatible processing algorithms. Let us briefly review the characteristics of these tools.

**Figure 1:**
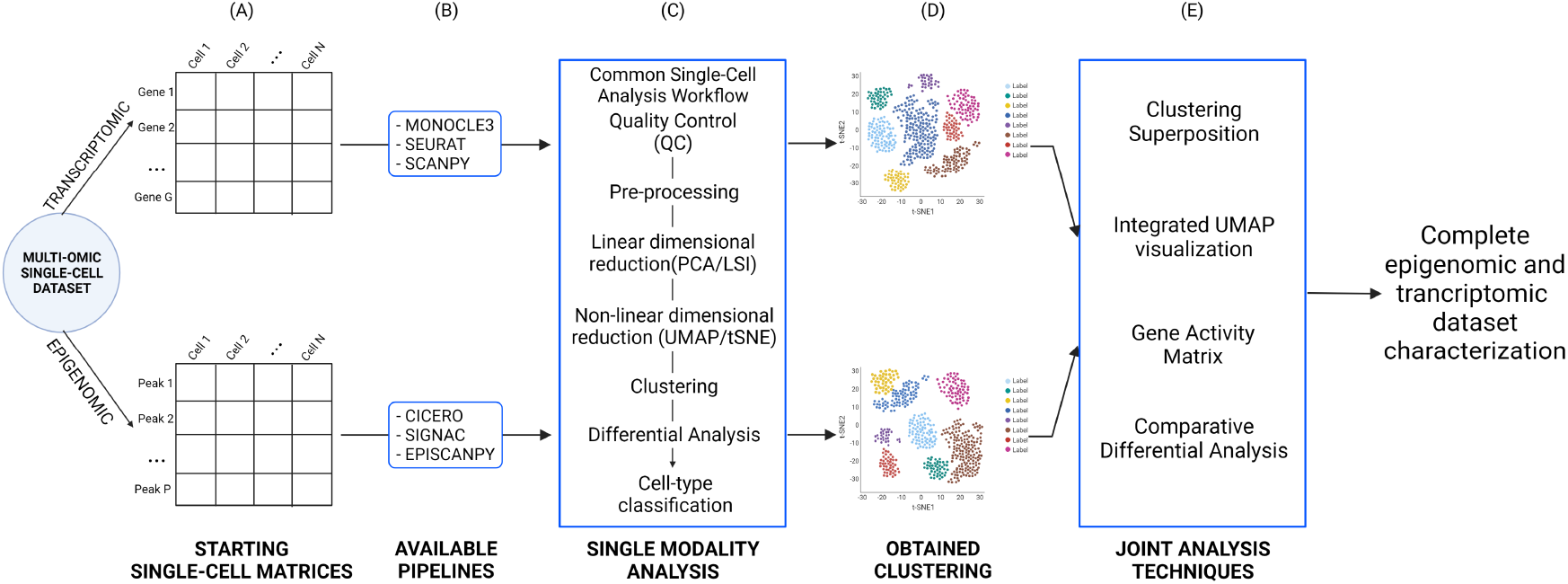
Schematic representation of the multi-omics analysis workflow. The pipeline receives the multiomics dataset, consisting of a scRNA-seq (G genes x N cells) and a scATAC-seq (P peaks x N cells) dataset (A). The workflow starts with the independent analysis of the datasets through the available pipelines (B), following the common single-cell analysis workflow (C). In this way, two separated clusterings of the same set of cells (D) are obtained. It then proposes a technique to integrate the two analyses (E) to obtain a complete epigenomic and transcriptomic dataset characterization.

Monocle3 is an R package for handling large scRNA-seq datasets. It supports data pre-processing, dimensionality reduction, and clustering, which are fundamental steps in single cell analysis [19]. Monocle3’s strength lies in its ability to perform single-cell trajectory analysis [35], which enables the characterization of cellular state transitions over time and is particularly useful for studying cell differentiation. Monocle3 is coupled with Cicero, a package that provides tools for analyzing scATAC-seq data. Notably, Cicero offers co-accessibility calculation [27], allowing investigation of genomic regions that tend to be simultaneously accessible. This feature provides insight into cis-regulatory interactions (e.g., enhancer-promoter interactions) and enhances our understanding of cellular regulatory mechanisms. Cicero follows a similar workflow to Monocle3 for processing scATAC-seq datasets.

Seurat [32] is an R package developed by Satija Lab [29] for quality control and analysis of scRNA-seq datasets. Seurat implements a well-organized data structure for storing datasets and research results, facilitating easy access and processing of available information. It provides functions for evaluating a set of quality control metrics commonly used in the community [14], ensuring the removal of lowquality samples that could negatively impact the analysis. Satija Lab also offers Signac, a companion package specifically designed for processing chromatin data (scATAC-seq). Signac employs the same data object as Seurat, enabling the storage of both data types and results in a unified structure, which is advantageous for studying multi-omics datasets. Signac provides specific quality control metrics [11] for scATAC-seq data and a workflow tailored to this data type. The Seurat and Signac combination offers a convenient way to store and process both scRNA-seq and scATAC-seq data uniformly, particularly useful for multi-omics analysis.

Scanpy is one of the few Python-based toolkits for single-cell gene expression analysis. It has gained popularity due to its flexibility compared to R-based tools. Scanpy offers all the necessary functions for data processing, similar to the pipelines mentioned above. One of its key features is the ability to handle large datasets (more than one million cells) [38] without requiring significant computational resources, which is particularly advantageous for scATAC-seq datasets. The companion package for Scanpy, Episcanpy, is a recently released toolkit for single-cell DNA methylation and scATAC-seq data. In addition to standard processing functionalities, Episcanpy allows users to create custom count matrices from raw data using any given set of genomic regions as features. This feature is handy for investigating the accessibility of specific genomic regions (such as enhancers and promoters) instead of typical peak-based approaches. Moreover, the scalability of the Python-based Scanpy package is particularly valuable when dealing with large and sparse scATAC-seq datasets [6].

This paper uses these three coupled pipelines to focus on the joint multi-omics analysis of scRNA-seq and scATAC-seq data. Regardless of the specific pipeline, the paper proposes a multi-omics analysis workflow described in Figure 1. It starts with an independent analysis of scRNA-seq and scATAC-seq datasets, followed by guidelines for integrating the two sets of results to use combined information to study cellular heterogeneity.

### 2.1 Single Modality Analysis

Regardless of the selected pipeline, the first step when performing a cell heterogeneity study using a multi-omics dataset is to analyze the different omics separately, obtaining independent results that can be compared. This activity, in turn, requires a deep understanding of the peculiarity of each dataset.

A scRNA-seq dataset can be represented as a sparse matrix, where the columns are associated with cells and the rows with genes (Figure 1 (A)). The matrix elements represent the number of RNA copies observed for a gene in a specific cell. The set of genes is consistent across different datasets for a given species, allowing easy comparison of results between experiments.

A scATAC-seq dataset is also represented as a sparse matrix, which is even sparser than scRNA-seq data. In this case, columns are associated with cells, while rows are associated with peaks. Peaks are intervals on the genome with a local enrichment of transposase cut-sites (Figure 1 (A)). Each peak is identified by its chromosome and genomic coordinate pair. The matrix elements in scATAC-seq data are binary numbers, where 1 indicates the accessibility of a genomic region (row) in a cell (column). The definition of accessible peaks in a dataset is derived from the experimental results, and therefore, peaks are not consistent across experiments as genes are in transcriptomic data. This characteristic makes the comparison of results from different experiments challenging.

Both datasets can be used to investigate cellular heterogeneity by following a similar workflow, including preprocessing, linear dimensionality reduction, non-linear dimensionality reduction, clustering, and differential analysis (Figure 1 (C)).

Pre-processing involves normalizing and scaling the data, which is necessary for all subsequent steps. For scRNA-seq data, Principal Component Analysis (PCA) is the preferred method for linear dimensionality reduction. In contrast, for scATAC-seq data, Linear Semantic Indexing (LSI) [25] is the typical approach that has shown the best performance on this type of data [31].

Two commonly used methods for non-linear dimensionality reduction and two-dimensional visualization of data are t-distributed Stochastic Neighbor Embedding (tSNE) [36] and Uniform Manifold Approximation and Projection (UMAP) [23]. UMAP is gaining popularity due to its ability to effectively separate different cells.

After applying linear and non-linear dimensionality reduction, the data are ready for cell clustering. Most pipelines implement clustering using Louvain [3, 16] or Leiden [34] clustering algorithms. The set of clusters represents the initial results obtained from any single-cell dataset, providing a data-driven classification of cellular heterogeneity.

Single-modality analysis creates two separate organizations of the same set of cells into clusters based on the multi-omics data (Figure 1 (D)). The features contributing to this organization and cell heterogeneity can be investigated through differential data analysis techniques.

Differential analysis is particularly effective for scRNA-seq data since the set of genes for an organism is consistent across different experiments. Cellular heterogeneity can be studied by examining specific marker genes whose expression characterizes and identifies particular cell types. Differential expression analysis performed on scRNA-seq data allows cell types to be labeled in a cluster based on previously known marker genes and discovering new markers [20].

Differential analysis of scATAC-seq results is more complex since these datasets do not have a consistent structure regarding features (rows). This characteristic makes it difficult to define the equivalent of a marker gene for accessibility data, and therefore assigning cell types based on Differential Accessibility (DA) data is not straightforward. One straightforward strategy is associating differentially accessible regions with the nearest gene, but this approach has limitations due to the simplistic assumption that proximity necessarily implies regulatory relations. Furthermore, the high sparsity of scATAC-seq data leads to lower-significance results, meaning that the differentially accessible regions are less representative of the clusters. Therefore, leveraging the respective transcriptomic information becomes crucial to overcome the lack of specificity in the accessibility features. This is one of the primary advantages of using multi-omics data. By employing gene expression information, one can investigate the cell type composition of a dataset and label the epigenomic part accordingly, creating a link between scATAC and scRNA data and enabling proper investigation of DA in cells.

Cell-type classification is the final step of this pipeline and is essential for studying cell heterogeneity through single-cell experiments. It can be implemented using two approaches. One approach involves analyzing the expression of marker genes, while the other uses labeling transfer techniques. The first approach investigates the expression levels of literature-based marker genes to label cells within a cluster. These markers are derived from various biological investigations that rely on protein-based assays, such as immunohistochemistry, to identify markers. Most biomarkers are thus proteomic, which poses some challenges in classifying transcriptomic data. Nevertheless, this is an easy and fast method that can provide at least general information about the dataset. To perform this type of classification, Monocle3 supports the integration of Garnett [26], an R package for automated cell-type classification. Garnett consists of a trained tissue/sample type-related classifier based on a given hierarchy with the related marker genes, allowing the application of the classifier to a dataset from the same tissue/sample type and obtaining a labeled dataset. The second approach is a relatively new technique that utilizes an externally labeled dataset and employs machine learning methods to predict the cell-type classification of the dataset of interest. This method heavily relies on the availability of a well-curated labeled dataset as a reference, which is only sometimes the case. Moreover, it is computationally more intensive than the previous approach. Seurat provides the Weighted-Nearest Neighbor (WNN) [32] analysis, directly integrated into its workflow.

Both approaches are viable for understanding the cellular composition of a dataset, at least at a high level, and are usually sufficient as a foundation for downstream studies.

### 2.2 Joint Analysis

The main goal of this section is to propose several ways to compare, connect, and jointly analyze scRNA-seq and scATAC-seq data pools, starting from their independent analysis (Figure 1 (E)). In particular, this article focuses on a new workflow that allows the superposition of clusters based on different omics and the Gene Activity Matrix (GAM) formalism, and the comparative analysis between differentially accessible and expressed genes identified by these different groupings. Given that Episcanpy is a relatively early pipeline, the possibilities for joint analysis are limited. Therefore, mainly Monocle3/Cicero and Seurat/Signac pipelines are used.

#### 2.2.1 Superposition and Joint visualization

As presented in the previous section, when processing single-cell datasets, one ends up with the cells visualized on a 2D space divided into clusters. This is true regardless of the omics considered and helps to have a first grasp of the cellular composition of the sample and, in general, the cell states identified with that particular biological information. Indeed, if the characteristics of the samples (i.e., species, tissues, region, status, etc.) are the same, such samples should support seampless joint analysis. Therefore, the results from scATAC-seq and scRNA-seq analyses performed on the same sample type should be directly comparable. Yet, due to technological limitations hampering reproducibility, comparing results from datasets originating separately is not trivial. On the other hand, multi-omics technologies perform analyses on multiple omics based on the same exact cells, ensuring intrinsic consistency between the omics analyzed, and making the subsequent analysis workflow more robust.

In particular, the first approach is to take the clustering labels from one analysis and superpose them on the other visualization. This way, one can intuitively highlight the differences between the results. For example, it can happen that, in one case, one group of cells is distinguished as two clusters, while in the other, it is only one, meaning that one omics is identifying some source of heterogeneity not detected by the other. Again, employing the coupled pipelines is helpful, since they store the results in the same object structures, making it easier to transfer the cluster labels.

As mentioned above, the cell-type classification employed by the gene expression data can be compared with the results of the epigenomic data. One has three different classifications of the same cells (i.e., transcriptomic clustering, epigenomic clustering, and cell-type labels) directly comparable to each other, allowing the investigation of the possible heterogeneity sources. To analytically evaluate how uniform the classifications are, it is advisable to employ the Adjust Rand Index (ARI) [15] and the Adjust Mutual Information (AMI) [37] from information theory, often used to compare clustering results. These metrics range between 1 and 0, where 1 is a perfect match while 0 is complete uncorrelation. To calculate these metrics, one can employ the R package aricode [9].

Moreover, one can visualize the cells in an embedded way, meaning a 2D visualization that considers both omics simultaneously. This is possible thanks to a Seurat asset, the WNN [32], an unsupervised framework enabling integrative visualization and downstream analysis of multi-omics datasets. This method receives the processed and dimensionally reduced data from the previous independent analysis and tries constructing a unique weighted graph, combining the two omics. One can further cluster cells by obtaining a clustering originating from both omics simultaneously. In this way, it is possible to exploit heterogeneity sources from gene expression and ATAC-seq, providing the most evident separation of the clusters.

#### 2.2.2 Gene Activity Matrix

Another way to link the information from these two different biological perspectives is through Gene Activity (GA) and, consequently, the concept of the GAM. GA refers to the general accessibility of a gene that allows transcription [27]. A GAM is, therefore, a matrix created starting from the accessibility data (in this case, scATAC-seq data), where the columns identify the cells, and the rows identify the genes. The matrix elements describe the accessibility of the gene in the cells, but this value is not univocally defined. It depends on the model chosen to interpret the relationship between accessibility and genes. All the pipelines mentioned above provide different ways to construct the GAM, mainly defining the accessibility of a gene through the accessibility of its promoter and gene body. These are overly simplistic models that describe the intricate correlation between gene regulation and gene expression, and a more thoughtful model would be valuable [21]. However, it is a good starting point that allows a middle ground between the biological levels, enabling a direct investigation of the correlation between epigenomic and transcriptomic information. The construction of the GAM by the pipelines is carried out in different ways, better explained and summarized in the respective papers [27, 33, 10], and also recapitulated in [21]. They start from the processed scATAC-seq data and obtain a matrix analogous to the general single-cell experiment datasets. From there, one can treat it like a gene expression matrix and process it, as discussed in the previous section.

In this way, the result is another processed dataset, where the features are the genes (comparable with the transcriptomic results), while its information content comes from the epigenomic data. Moreover, one can compare the resulting clustering with the previous ones, adding another layer of information.

#### 2.2.3 Comparative Differential Analysis

The final part of the joint study focuses on a comparative differential analysis, which utilizes the newly obtained Gene Activity Matrix (GAM) to compare genes on their transcriptomic and epigenomic levels. The emphasis is placed on informative genes, such as differentially expressed genes or cell-type markers. The goal is to examine the accessibility of these features, using the concept of activity, and understand their non-trivial correlation with gene expression. This approach determines if the transcriptomic features that characterize a particular cell type are also reflected at the epigenomic level.

To begin the analysis, the processed GAM with its associated clustering classification is used. A differential activity analysis results in a list of genes whose activity characterizes the identified clusters. This allows one to identify specific genes that are differentially active and differentially expressed, contributing significantly to the heterogeneity of biological information. However, it is not immediately apparent whether these genes characterize the same cells, as the clustering obtained from the GAM may not be as accurate and distinct as previous clustering methods. Another round of differential activity analysis is conducted separately for the clustering obtained from the processed gene expression data clustering, along with cell-type classification. This ensures that the obtained results, specifically the differentially active genes, are directly related to the clusters identified in previous analyses. Consequently, they can now be compared with the differential features observed within the same groups of cells.

In conclusion, comparative analysis facilitates exploring the intricate relationship between the accessibility and expression of genes related to cell types. Additionally, validating the differential activity of marker genes for the same cell types helps establish the credibility of the dataset’s classification.

## 3 RESULTS

This section illustrates the application of the presented multi-omics analysis workflow on a 10k human PBMC multi-omics dataset from 10XGenomics [30, 1]. The dataset comprises 10,691 human peripheral blood mononuclear cells. The purpose is to showcase the obtained outcomes and guide their interpretation. To streamline the presentation of results, we solely include the key findings derived from the bioinformatic pipelines employed. However, comprehensive results for each employed pipeline, accompanied by dedicated and detailed explanations in code notebooks, can be accessed on the online repository at [Zenodo].

### 3.1 Single Modality Analysis

The scRNA-seq analysis produces the first unsupervised clustering of cells, as illustrated in Figure 2. The selection of resolution parameters adheres to recommended guidelines, with values falling within the typical range for datasets of this scale. The figure exhibits all three UMAP embeddings and the clusters generated within each pipeline.

**Figure 2:**
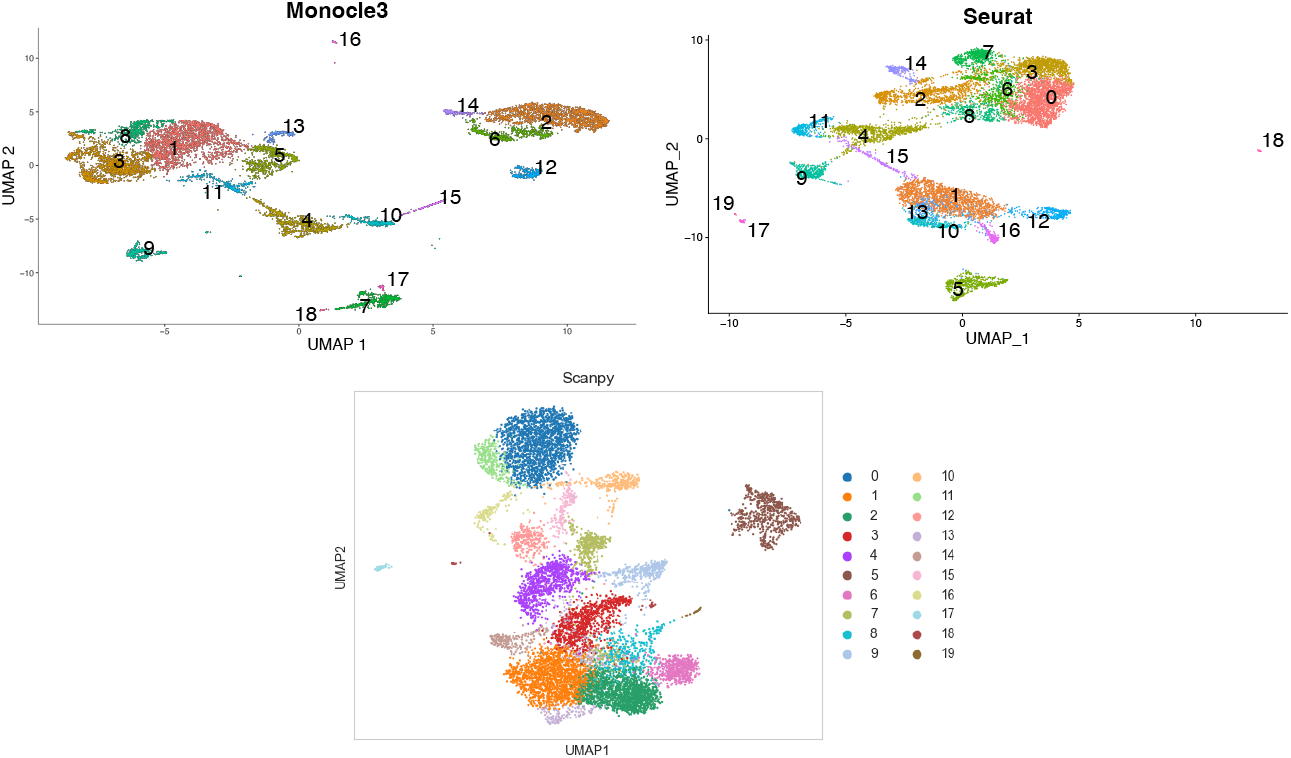
2D UMAP visualization of the dataset obtained from scRNA-seq data with Monocle3 (left), Seurat (center), and Scanpy (right) pipelines. The clustering algorithms led to about 20 clusters. In general, one can see two bigger populations, two smaller ones, and some isolated few-cell populations. This is in line with what we expect from a PBMC datasets, even if, at this point, the actual cell types are unknown.

Observing Figure 2, it becomes evident that cells tend to aggregate into three distinct populations, which probably correspond to T cells, Monocytes, and B cells. Across all instances, the total number of clusters is approximately 20. However, at this stage, the specific cell types remain uncertain. The next step involves conducting a Differential Expression (DE) analysis to address this. Supplementary material Figure S1 displays the DE analysis results, highlighting known genes associated with particular cell types (e.g., FCGR3A for Monocyte clusters).

The scATAC-seq analysis also involves an independent and unsupervised clustering of the cells (Figure 3). Due to the characteristics of accessibility data, the resulting clustering tends to be less precise and well-defined compared to the previous analysis. This is particularly evident in the case of Episcanpy (Figure 3, right). Additionally, setting a higher resolution parameter to achieve the same number of clusters is generally necessary. In general, accessibility data alone are apparent to be less suitable for studying cellular heterogeneity compared to gene expression data. As discussed above, this observation is further supported by the DA analysis of the peaks (Supplementary Figure S2), which fails to provide a precise cluster characterization.

**Figure 3:**
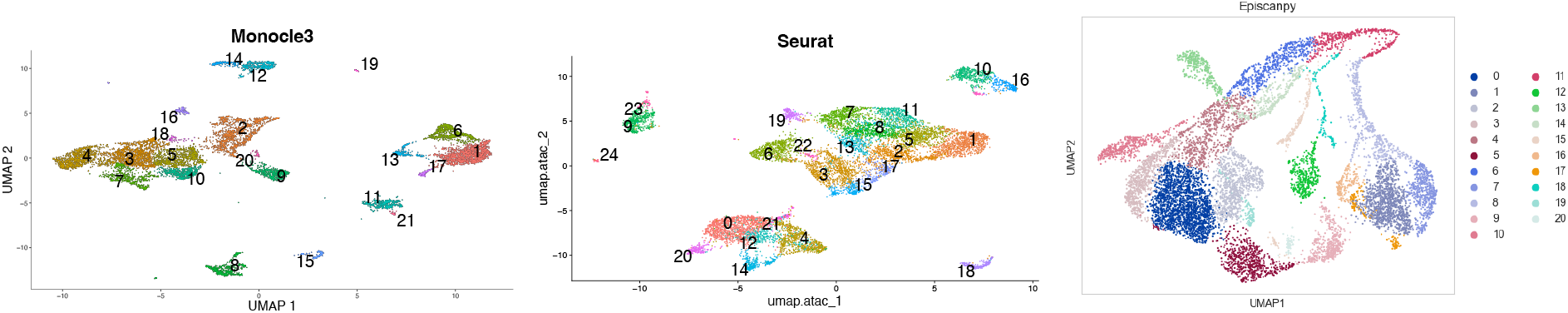
2D UMAP visualization of the dataset obtained from scATAC-seq data with Monocle3 (left), Seurat (center), and Scanpy (right) pipeline. The resolution of 0.8 led again to 19 clusters. In general, two larger populations, three smaller ones, and some isolated few-cell populations can be seen. This aligns with what we expect from PBMC datasets, even if the actual cell types are unknown. It seems that cells, in this case, tend to be more aggregated, showing the differences and limitations of study cellular heterogeneity with scATAC-seq data concerning scRNA-seq.

However, the subsequent step involves the proper classification of cells using two different methods which are described below. Firstly, the Garnett method offers convenient integration with Monocle3. In this case, a pretrained PBMC classifier provided by Garnett is used. The implementation of classification is straightforward and results in near-complete labeling of the datasets (Figure 4). The classification is performed at a coarse level, dividing cells into a few major cell types, which helps to identify the cell type of the primary cell groups, but does not capture the subtypes. This limitation arises from the inherent nature of the chosen classifier, which operates at this resolution level. Figure 4 shows some cells as “Unknown,” meaning that, according to Garnett’s classification, they cannot be identified confidently.

**Figure 4:**
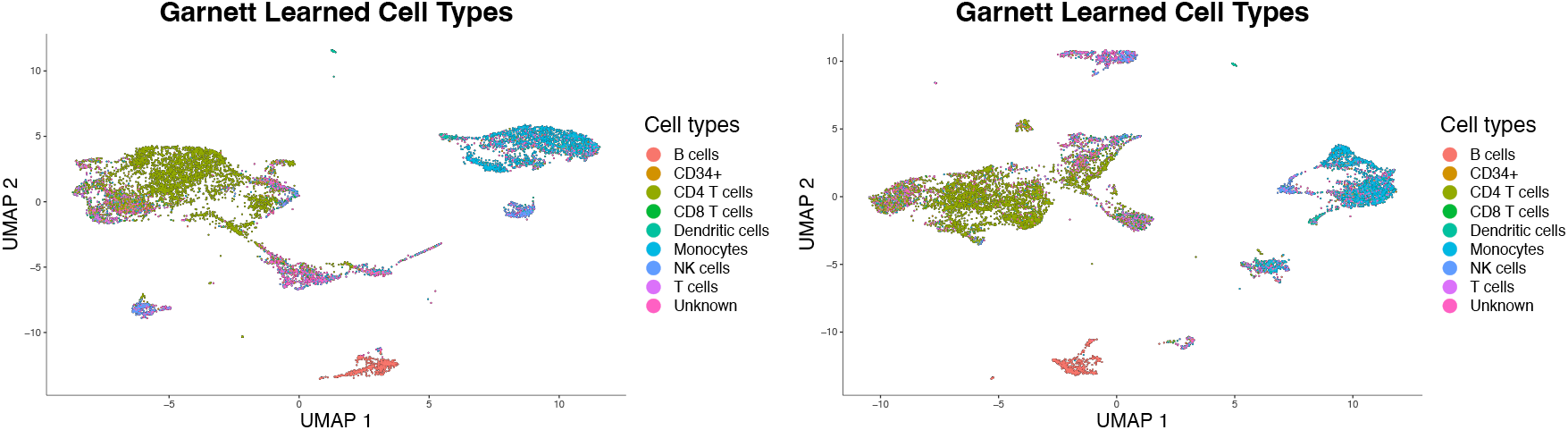
Cell-type labels from Garnett on scRNA-seq (left) and scATAC-seq (right) Monocle3 visualization. A total of 8 cell types are present (plus one Unknown class). The biggest population composes of T cells (green and purple), with near them the Natural Killer (NK) cells (blue). The second one represents Monocytes and Dendritic cells (light blue). The smaller groups consist of B cells (red).

The second method involves label transfer, facilitated by Seurat. In this case, a curated and processed SeuratObject of a labeled dataset needs to be loaded. The level of detail achieved depends on the selected reference dataset. Multiple taxonomies are available, but here we present the methodology for one of them. The results in Figure 5 demonstrate that the labels obtained through label transfer align with those obtained by Garnett, but with a more detailed taxonomy.

**Figure 5:**
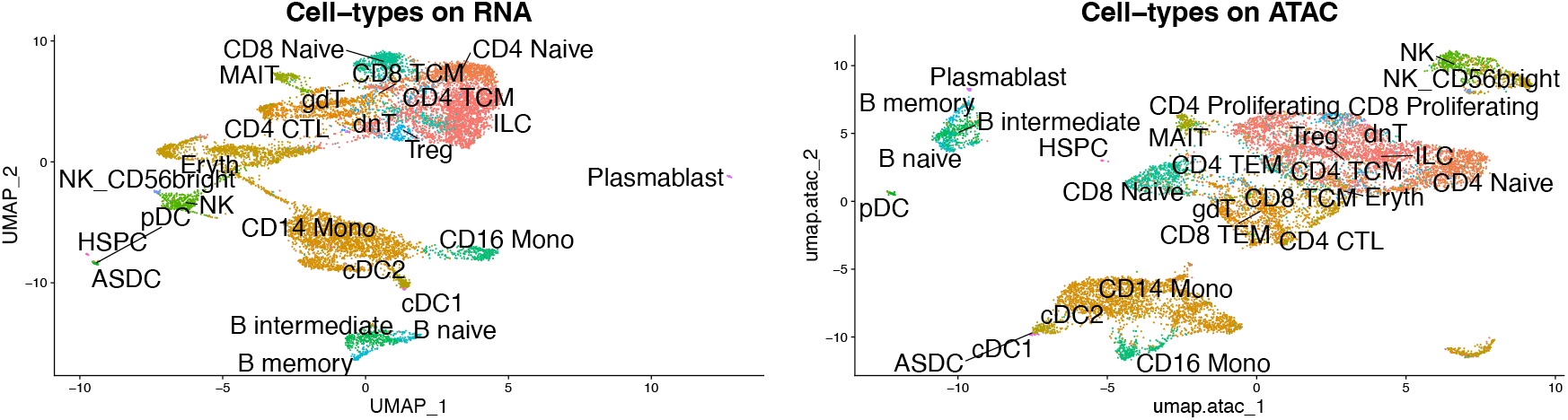
Cell-type labels from Seurat integration on scRNA-seq (left) and scATAC-seq (right) Seurat visualization. A total of 28 cell types are present. The biggest population comprises T cells divided into many subtypes (mainly between CD4 and CD8 T cells) with the Natural Killer (NK) cells near them. The second one represents Monocytes and Dendritic cells. The smaller groups consist of B cells, divided into three subtypes. Isolated groups of a few cells are particular and rare cell types. It is worth noticing the different separation of some groups of cells, like the NK cells, which appear more homogeneously separated in the scRNA-seq case.

Both methods yield a classification of cells based on gene expression data. Additionally, due to the multiomic nature of the datasets, the cell type composition of clusters obtained from the scATAC-seq analysis can be readily determined (Figure 4, Figure 5).

The results obtained align with the expected cell types. The largest population consists of T cells, further divided into subtypes primarily between CD4 and CD8 cells, with Natural Killer (NK) cells located nearby. The second most populous group comprises Monocytes and Dendritic cells. B cells are also present, with three subtypes identified. Some cell groups, such as Plasmablast cells, also represent rare cell types. Some interesting observations can be made from the scATAC-seq data results (Figure 5, right). Specifically, CD4 and CD8 cells exhibit clearer subdivisions within the larger population, and the separation of NK cells is enhanced compared to previous findings. It is intriguing to note that in this case Plasmablast cells are located near B cells, suggesting their origin from the B cell population.

However, the results of the scATAC-seq data present specific challenges. For example, the subdivision between subtypes tends to be less precise, with subtypes displaying greater mixing. Moreover, a small population of cells remains unidentified and consists of different cell types.

This initial step enables data exploration based on biological information, forming subsequent analyses’ focus.

### 3.2 Joint Analysis

The joint analysis aims to establish connections between the results obtained from the individual omics analyses. Due to the multi-omics nature of the datasets, transferring metadata associated with one omics to another becomes straightforward, as the cell IDs remain consistent and no integration is required. Consequently, it is natural to compare the clustering results by superposing the cluster labels on the 2D visualizations, as depicted in Figure 6 using Monocle3 (left) and Seurat (right).

**Figure 6:**
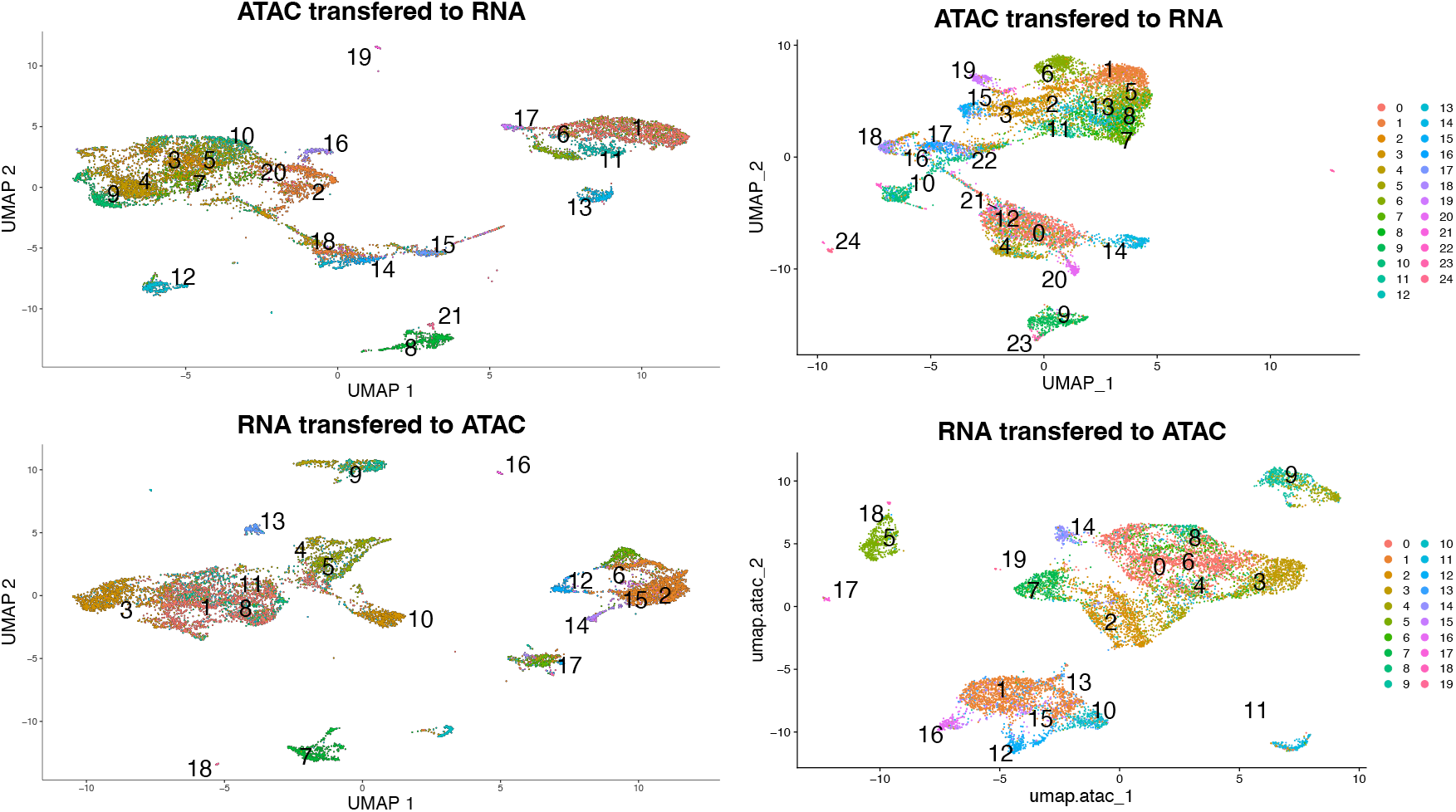
Superposition of cluster labels on the other omics 2D visualization in Seurat. By doing so, one can investigate how consistent the cluster results are between the two separate omics analyses. There is a clear consistency between the clusters, which means that the clusters have a clear correspondence with the other omics. The most uncertainty comes from the subtypes within the main population, which, especially for the scATAC-seq case, tend to be very intermingled.

These observations demonstrate that, in general, the clustering results exhibit consistency across omics, albeit without achieving a perfect match. Nevertheless, a general correspondence is evident, indicating that the analysis performed on the different datasets identifies similar sources of heterogeneity. However, as discussed earlier, notable differences persist, particularly regarding the extent of separation between specific subtypes within the same populations. This observation suggests the presence of omics-specific sources of heterogeneity.

To address this, the WNN analysis, exclusively provided by Seurat, can generate a unified 2D visualization of the dataset, incorporating information from both omics. Figure 7 depicts the results of this analysis. By combining the omics, this representation captures discernible features, such as the distinction between Dendritic cells (more prominent in scRNA-seq data) and the clearer separation between CD4 and CD8 cells (more evident in the scATAC-seq data). The significance of this analysis lies in the initial exploration of this type of data often relies on visual inspection using UMAP plots. Therefore, having this integrated view facilitates a comprehensive assessment of the heterogeneity of the dataset.

**Figure 7:**
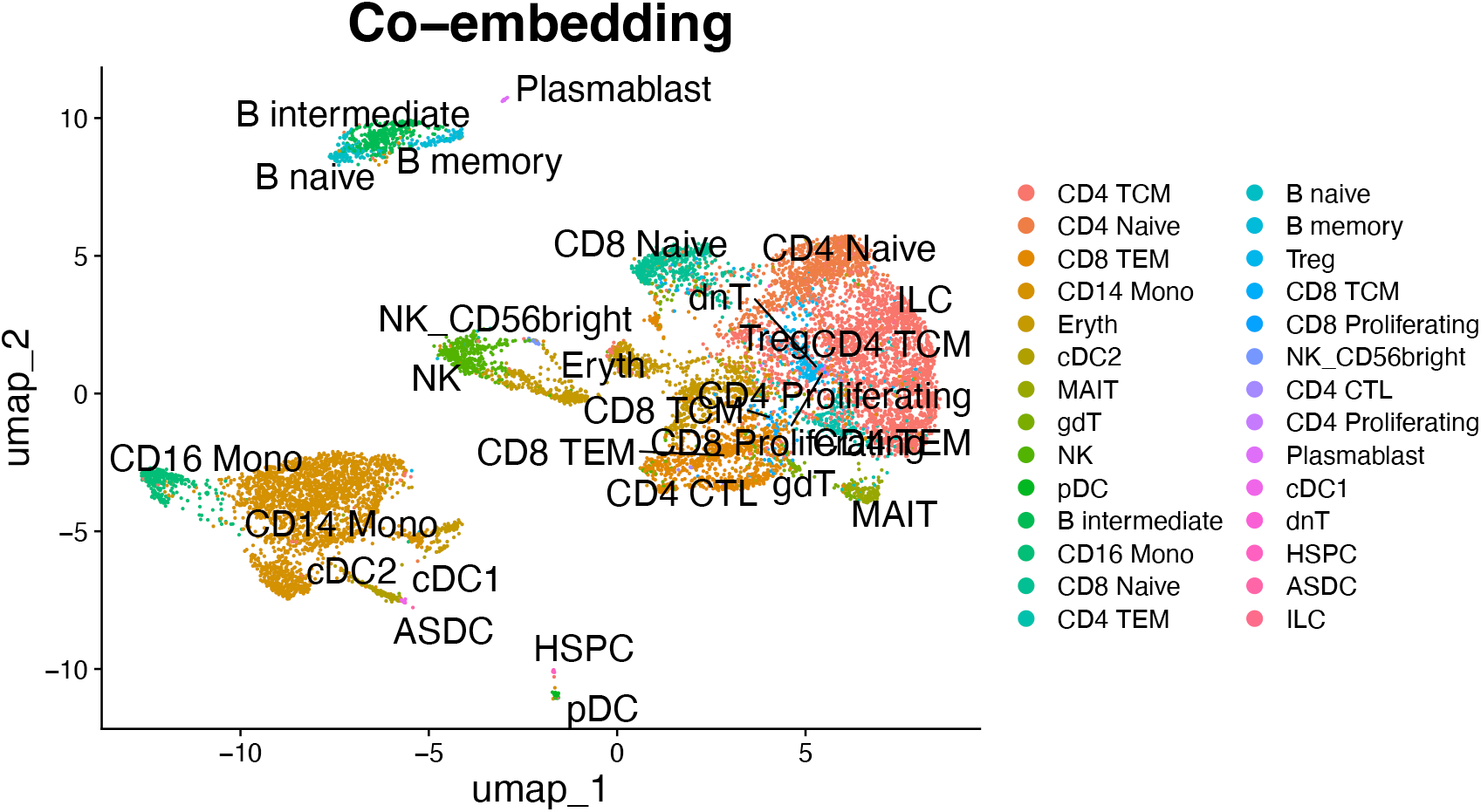
Integrated visualization of cells based on the Weighted Nearest Neighbor Analysis from Seurat. This allows the construction of a graph that integrates the omics and defines cell states based on both omics. In this way, one obtains a visualization that synthesizes what is obtained from the separate omics, obtaining a better separation between cell states. For example, there is better separation between CD8 and CD4 T cells (more evident in the scATAC-seq data), and the Dendritic cell subtypes and the Monocytes are more separated within one another (this is more evident in the scRNA-seq data).

The subsequent step involves generating the GAMs using the methods proposed by Cicero (resulting in a matrix comprising 15,608 genes) and Signac (resulting in a matrix containing 19,607 genes). As mentioned, the GAMs are datasets processed similarly to the scRNA-seq data. The results shown in Figure 8 are already annotated with the cell types obtained previously. Given the significant manipulation of the data involved in the GAM construction, studying these results alone does not provide novel and specific conclusions, particularly regarding heterogeneity. However, the composition of the clusters remains consistent with previous investigations. However, it can be concluded that exploring the GAMs primarily enables the detection of differences between broad cell types (e.g., T cells and B cells) rather than finer distinctions between subtypes. However, the GAM provides a direct and interpretable link between gene accessibility and expression. Particularly intriguing is the exploration of marker genes and their corresponding accessibility patterns. The results of both Signac and Cicero reveal the accessibility profiles of genes such as CD8A, MS4A1 and CD14 (Figure 9), which align with their expected associations of cell types (i.e., CD8 T cells, B cells, and Monocytes, respectively). This is an illustrative example of the potential to study the nontrivial relationship between gene accessibility and expression.

**Figure 8:**
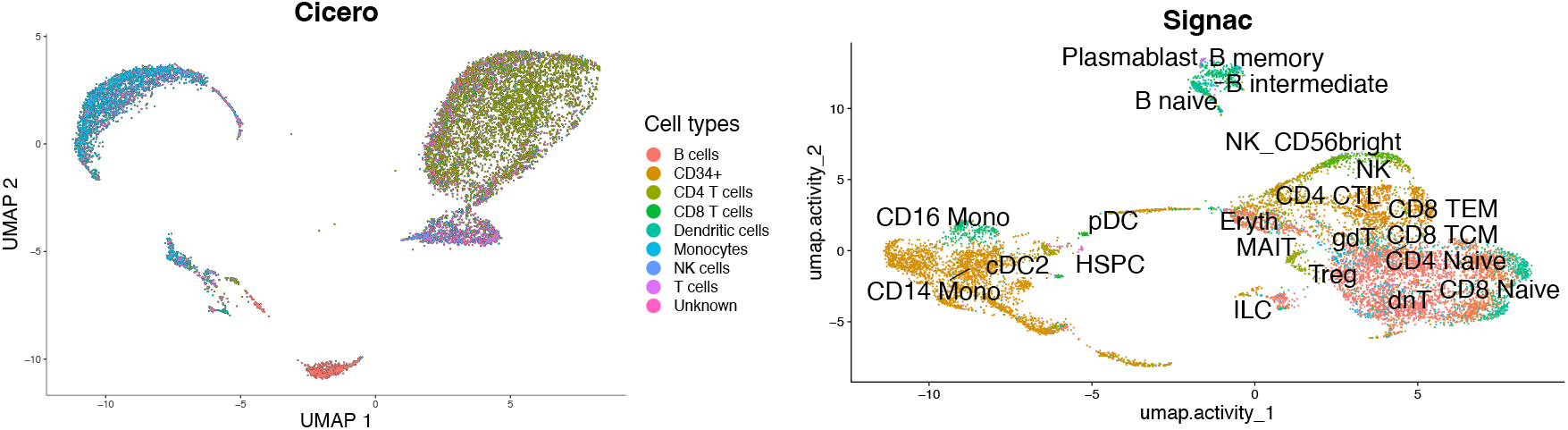
Cicero (left) and Signac (right) GAM results, labeled with the respective method’s cell types. Again, there are two main groups of cells, plus some smaller populations. In general, with the GAM, the cells tend to be less separated, and it is not optimal for a cell state investigation, with many subtypes mixing, especially in the Signac case.

**Figure 9:**
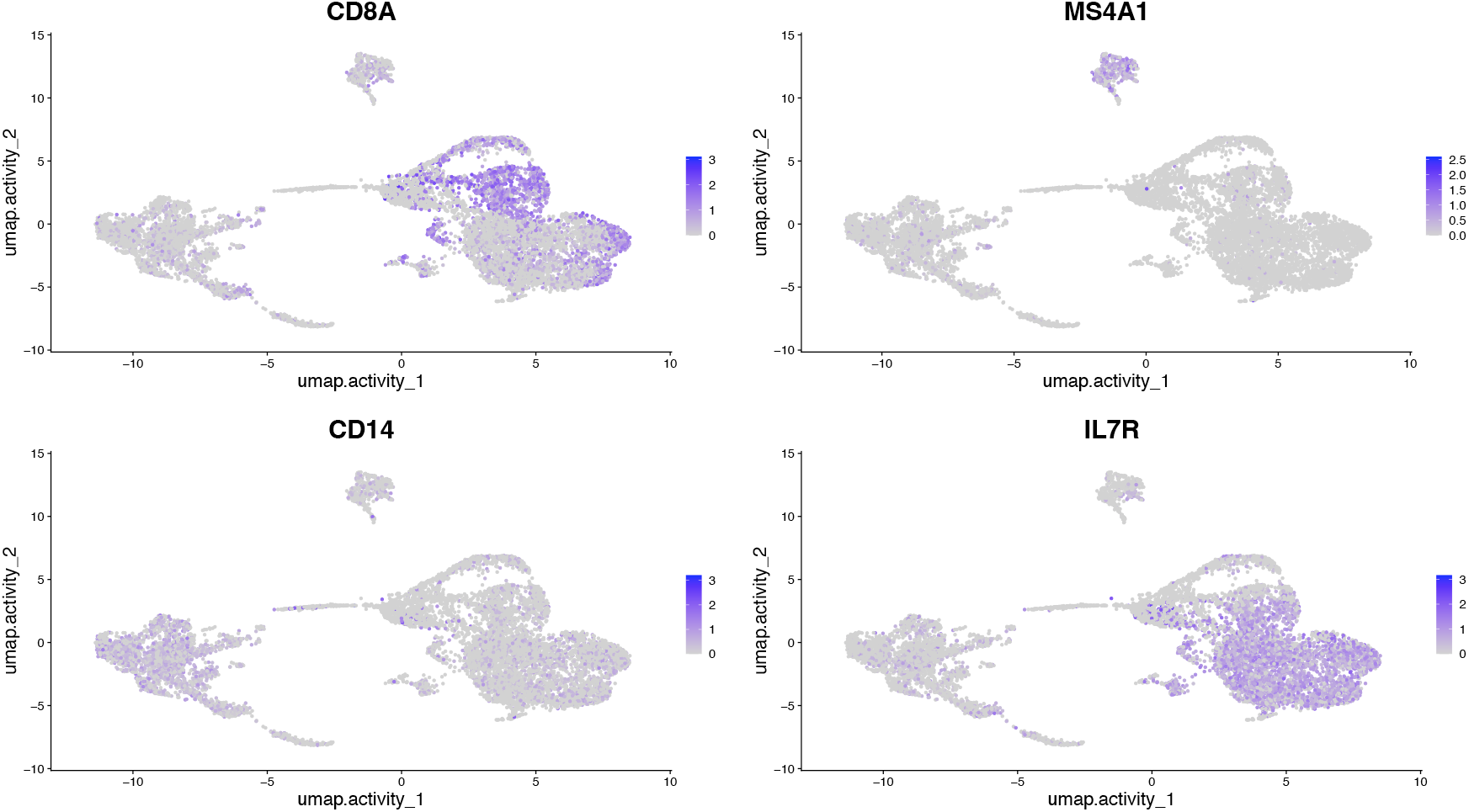
Activity visualization of some genes. They appear in the expected cell types: CD8A is active in respective CD8 T cells, MS4A1 is active strictly in B cells, CD14 is active strictly in monocytes, and IL7R is active mainly in CD4 T cells.

From this point, performing differential analysis, specifically comparative differential analysis naturally follows. Initially, differential analysis conducted on the GAM uncovers genes that exhibit differential accessibility in specific cell types, which may not have been evident from differential expression analysis alone. This highlights the ability of the GAM approach to identify features that characterize cell types, thus defining markers from the perspective of gene accessibility.

The final step in the joint analysis assesses whether the genes characterizing cells are consistent across both biological levels. To achieve this, differential analysis is conducted on the GAM, considering different partitions such as cell types and scRNA-seq clusters and examining the resulting genes. The results obtained using Monocle3 indicate that CD8A, IL7R, and MS4A1 exhibit differential accessibility in the same cell populations, irrespective of the omics (cell types or scRNA-seq clusters). These genes serve as robust markers for multi-omics datasets, as they can effectively distinguish cell populations regardless of the omics employed. Consequently, it is advisable to study this new class of markers, characterized by their consistent behavior in comparative differential analysis.

In conclusion, the GAM facilitates the integration of the two biological levels and aids in understanding the intricate correlation between epigenetic modifications and gene expression regulation. Furthermore, it enables the development of models to study this relationship in greater detail.

## 4 CONCLUSION

This study presents a comprehensive bioinformatic analysis of multi-omic scRNA-seq and scATAC-seq data. The focus is on identifying optimal pipelines for separate and joint analysis of these datasets. Seurat, Monocle3, and Scanpy, along with their associated packages Signac, Cicero, and Episcanpy, are effective pipelines for multi-omics analysis due to their dedicated packages for each omics. The separate analysis follows the standard workflow for single-cell data analysis, resulting in a 2D visualization of cell clusters based on different omis. Cell type labels are assigned to understand the cellular heterogeneity of the dataset, utilizing Seurat’s WNN and the Garnett package linked to Cicero. Since they represent the same cells, cell-type labeling is applied to gene expression data and scATAC-seq clusters.

The most intriguing aspect lies in the integration and joint analysis of the two omics. As the clustering results are directly comparable between omics, it becomes possible to investigate differences in the representation of cellular heterogeneity and clustering between them. Additionally, Seurat’s WNN facilitates the integration of information from both omics into a single 2D visualization, leveraging the strengths of both data types.

Furthermore, the concept of GAM is crucial in linking different features, such as peaks with their accessibility and genes with their expression. The GAM provides insights into the overall accessibility of genes, enabling the identification of genes that characterize cell types from a different perspective. Comparative differential analysis yields two notable results. Firstly, it examines whether marker genes (identified based on expression) and differentially expressed genes are also differentially active in the same cell groups, highlighting their reliability across multiple omics. Secondly, differential activity analysis uncovers a novel class of epigenomic marker genes that may not be identified solely from transcriptomic data.

In conclusion, this study aims to establish best practices for analyzing multi-omics scRNA-seq and scATAC-seq datasets. These datasets provide a unique opportunity to gain deeper insights into cellular states and require a comprehensive understanding of the analyzing of different omics together.

## 5 DATA AVAILABILITY

The 10X dataset (10Xmulti) was derived from sources in the public domain: https://www.10xgenomics.com/resources/datasets/10k-human-pbm-cs-multiome-v-1-0-chromiumcontroller-1-standard-2-0-0. The code underlying this article is available in Zenodo, at https://doi.org/10.5281/zenodo.8188656.

